# Structural signatures of salinity adaptation in *Microviricetes* capsids

**DOI:** 10.64898/2026.11.10.765580

**Authors:** Varada Milind Khot, Mart Krupovic, Bas E. Dutilh

## Abstract

Phages, like their bacterial hosts, have evolved to proliferate across the full range of environments on Earth, but the specific amino acid residues that underlie their environmental adaptations remain elusive. Here, we developed a computational framework that uses structure-based multiple sequence alignments, a random forest classifier, and a phylogeny-aware regression model to find amino acid residues linked to specific environments.

Applying this framework to public metagenomic data, we analyzed 1,110 *Microviricetes* major capsid proteins (MCPs) from the lake and ocean biomes, and found that ocean virus capsids were highly enriched in negatively charged residues on internal and external surfaces. This adaptation likely helps maintain solubility at high salt concentrations and stabilizes ssDNA-capsid interactions by sequestering positively charged ions inside the capsid. At intersubunit interfaces, ocean capsids showed enrichment of cation-π and π-π interactions, which are more resistant to solvation than salt bridges and hydrogen bonds. Amino acid residues in lake capsids were more often substituted by biochemically dissimilar amino acids in ocean capsids than the reverse.

Together, these results show that microvirus capsids adapt to high-salt environments through changes in surface charge and substitutions affecting intersubunit interactions. Further, we describe a general approach for studying the adaptation and stability of any protein family under complementary environmental or phenotypic conditions.

## Introduction

Microbes and their viruses adapt continually, not only to one another but also to the external environment in which they persist. Bacteriophages are found in virtually every habitat on Earth, and their proteinaceous capsids are directly exposed to the surrounding medium. This strong environmental selection, combined with a small set of conserved major capsid protein folds^1^, makes phage capsids well suited to comparative analyses of structural stability. How these natural assemblies remain intact under environmental stress is also of practical interest, as it offers design principles for engineering more robust capsids. Such optimization is already established for adeno associated viruses (AAVs), which are widely used as vectors for drug and vaccine delivery^2,3^. As the application of phage therapeutics grows, comparable efforts will be needed to make phage capsids and other proteins resistant to varying conditions. Naturally stable capsids offer a direct source of principles to build upon.

Recent comparative genomic and structural studies have begun to explore how phages stabilize their capsids^4^, including under challenging environmental conditions such as the high temperatures of hot springs^5^. These studies point to intersubunit interactions stabilized by isopeptide bonds and extended loops, which allow the capsid to remain flexible. An important open question is whether similar patterns contribute to capsid structural stability across other environmental gradients, such as pH or salinity. Salinity represents a compelling axis along which to investigate capsid adaptation, as it is one of the main dimensions structuring the microbial world^6^. Ions in saline water may readily interact with surface-exposed residues, interfere with protein-protein interactions at subunit interfaces, and alter viral adsorption to host cells^7^. In high-salinity environments such as marine ecosystems, capsids must maintain structural integrity against osmotic stress and reduced water activity, as water is drawn away from the protein toward charged ions in solution^8,9^. Most studies of protein haloadaptation in both cells and viruses report an enrichment of acidic residues on the external protein surface that bind hydrated cations^9–14^.

We explore the mechanisms of capsid stability across the class *Microviricetes* (formerly known as the family *Microviridae*) within the realm *Volvereviria*^15^. Originally described through the archetypal phage φX174, many *Microviricetes* members have since been recovered by metagenomic sequencing, revealing diverse lineages^16,17^ that colonise environments ranging from the mammalian gut to the open ocean^18^. Microviruses are non tailed bacteriophages encoding a compact single stranded DNA genome of 4.5 - 6 kb and assembling an icosahedral capsid with T=1 geometry^17^. Their capsids are built from major capsid proteins (MCPs) with a single jelly roll fold, making microvirus MCPs a tractable model for how viral structural proteins adapt to different environments. Indeed, a recent study tracking microvirus lineages from sewage treatment to river water showed that environmental factors were the strongest determinant of microvirus populations^19^.

We ask whether capsids of microviruses from freshwater and marine ecosystems carry an identifiable signal of adaptation to the respective environments and position the adaptive changes in amino acid sequence in the context of the three dimensional structures of the microvirus capsids. We mined public metagenomes for *Microviridae* MCPs from lake and ocean biomes and developed a generalizable pipeline that identifies biome associated residues from protein sequences and metadata alone, by combining a random forest model with a phylogenetically aware generalized linear model to account for shared viral ancestry. Mapping the resulting residues onto a reference capsid allowed us to interpret the changes in the context of the capsid structure, linking comparative genomics to structural biology and viral ecology.

## Results

### The global phage capsid protein space

To capture the diversity of phage capsids, we surveyed public metagenomes from the IMG/VR^20^ and MGnify^21^ databases, which yielded 1.4 million globally dispersed major capsid proteins (MCPs) from aquatic biomes (Fig. 1A). We selected the 3,000 viral capsid protein sequences annotated as Prokaryotic Virus Remote Homologous Group PHROG_514, which contains the *Microviricetes* capsids^24^. A sequence similarity network showed that proteins have many more connections within (lake-lake, ocean-ocean, n=3,410) than between biomes (lake-ocean, n=245, Fig. 1B).

**Figure 1.**
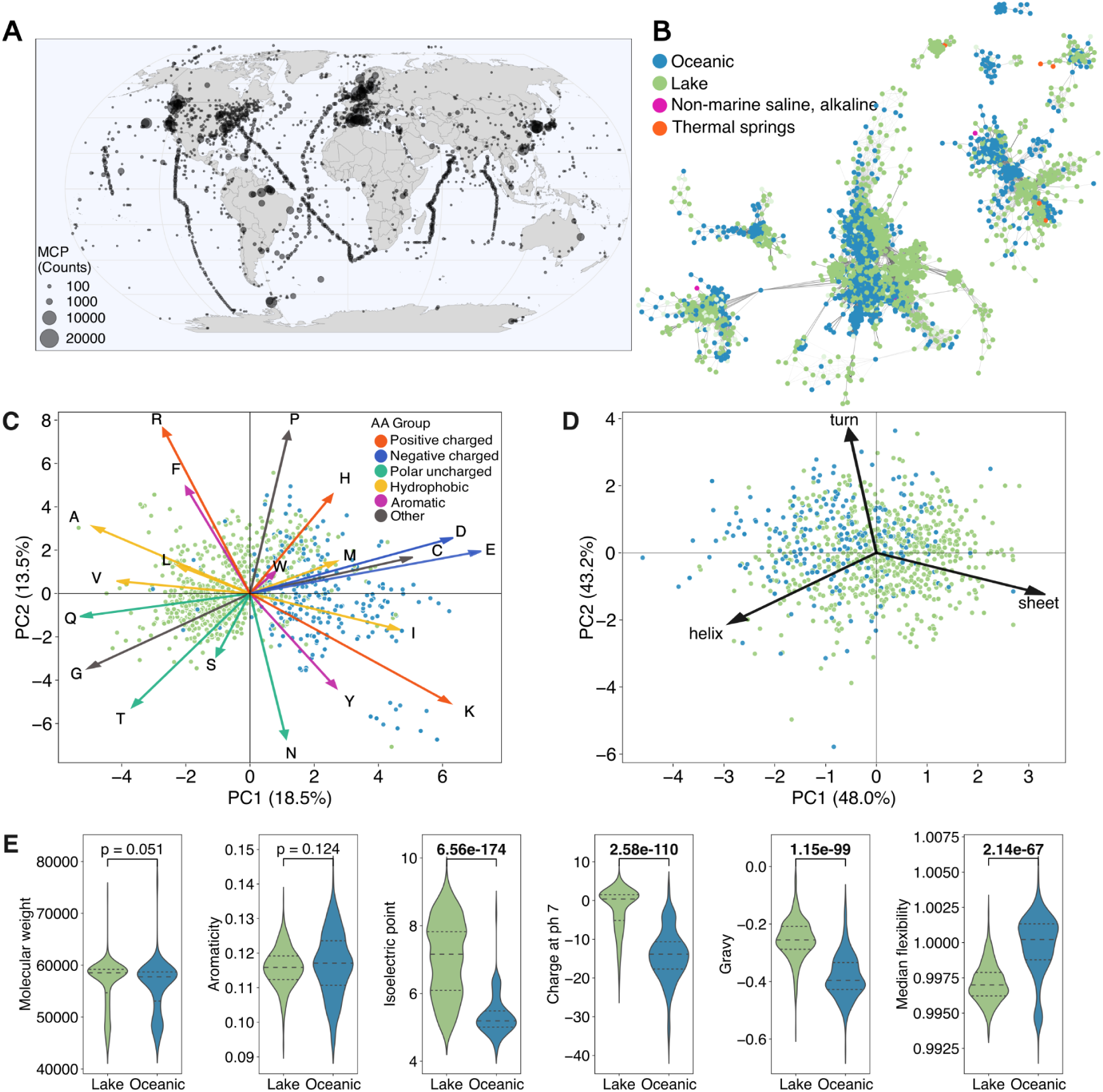
A) Locations of origin of major capsid proteins found in IMG/VR ^20^ and MGnify ^21^ metagenomes (n=1.4 M). B) Sequence similarity network (Diamond BLASTp ^22^, evalue < 10^-5^) of Microviridae major capsid proteins, coloured by biome of origin. Only connected components with more than 25 members are shown. C) PCA of amino acid vectors and (D) vectors of predicted secondary structural elements from “Lake” and “Oceanic” capsid protein sequences (n=824 and 286, respectively). E) Physicochemical properties of “Lake” and “Oceanic” capsid proteins calculated by Biopython v1.6^23^. P-values were generated using Welch’s t-test to test whether the differences between the two biomes were significant. The distribution of protein properties along the alignment length is shown in Figure S1.

Despite all sharing a T=1 capsid geometry, microviruses can have substantial morphological differences. For example, the best-known microvirus, ɸX174 (order: *Bullavirales*), forms capsids with the pentameric receptor-binding spike complexes located at the 5-fold axes of symmetry^25,26^, while members of most other lineages of *Microviricetes* lack the spike protein and instead were shown or predicted to contain an insertion domain within the MCP which forms receptor-binding protrusions at the 3-fold axes^27–30^. To ensure comparability between our proteins for downstream analyses, we selected only MCPs with structural homology to the *Gokushovirales* MCP sequences, resulting in 1,110 capsid protein sequences (Fig. S3) with lengths between 400-600 amino acids. These were clustered together based on structural homology using Foldseek^31^. The removed sequences are further indicated in Figure S3 by red dashed lines.

Next, we investigated the composition of these lake and ocean protein sequences. A principal component analysis (PCA) of amino acid frequency vectors showed a clear differentiation between lake and oceanic *Microviricetes* MCPs along the first principal component (Fig. 1C). This separation was driven primarily by an enrichment of aspartate (Asp, D), glutamate (Glu, E), cysteine (Cys, C), isoleucine (Ile, I), and lysine (Lys, K) in oceanic proteins, and arginine (Arg, R), phenylalanine (Phe, F), alanine (Ala, A), leucine (Leu, L), valine (Val, V), glutamine (Gln, Q), glycine (Gly, G), and threonine (Thr, T) in lake proteins. An enrichment in Asp and Glu is consistent with amino acids found in halophilic bacterial and archaeal proteins, suggesting this as a global adaptation to salt^9–11,32,33^. In contrast, Lys has been observed to decrease in halophilic adaptation of proteins^33^.

Lake and ocean capsids also differed significantly in their secondary structure content, isoelectric points, net charges at pH 7, hydropathy index (GRAVY: grand average of hydropathy) and median flexibility. With a lower isoelectric point (median: pH 5.25) and a median net charge at pH 7 of -15 e, oceanic capsid proteins were predominantly negatively charged. In contrast, lake proteins were neutrally charged, with a wider distribution of isoelectric points (median: pH 7) and a median charge at pH 7 of 0 e. Oceanic proteins were also enriched in hydrophilic residues and had flexible side chains, as they had a lower GRAVY index and slightly higher Vihinen flexibility scores^34^ than lake proteins. The lower flexibility of lake proteins may be partly explained by their higher β-sheet content (Fig. 1D), rigid secondary structures that consist of a planar hydrogen-bonding network^35^. The differences in flexibility were concentrated between positions 70-120 and in the receptor binding domain region, while charge differences were distributed along the protein length (Fig. S1). Microvirus genome lengths of capsids used in this study also differed between biomes, with microviruses from a lake environment with slightly larger genome lengths than those from an ocean environment (Fig. S2). We hypothesize that the enrichment of negatively charged, hydrophilic, and flexible amino acid residues in oceanic *Microviricetes* capsids reflects an adaptation to the high-ionic-strength marine environment where electrostatic and solvent interactions may be critical for capsid stability.

### Random forest and phylogenetic regression analyses identify lake- or ocean-associated residues

To identify which specific amino acid residues, and at what positions in the protein structure, drove the separation of MCPs from lake and ocean environments, we developed a workflow to associate individual residues with each biome (Fig. 2A). First, all protein sequences were aligned to a reference sequence (ɸEC6098, purple star in Fig. 2B) to generate a multiple sequence alignment. Amino acid-based multiple sequence alignments have been long used to compare proteins, find mutations and build phylogenetic trees as they hold key information about gene evolution. They have also been previously used to predict functional changes in protein subfamilies^36^. Here, we used a one-hot-encoded version of our multiple sequence alignment to define features, each corresponding to a specific position and amino acid combination (Fig. 2C), which served as the input to a random forest (RF) model trained to predict the biome of origin for each sequence. The RF model achieved a balanced accuracy of 93%, indicating that a strong environmental signal was present in the amino acid sequences, consistent with the PCA separation described above. Ranking features by Gini score further allowed us to extract those that contributed most to the separation between ocean and lake capsid sequences. A caveat to the RF model is that it cannot distinguish between residues that are genuinely biome-associated and those shared among phylogenetically related viruses that appear in the same environment. To account for this phylogenetic bias, we used phyloglm^37^ to identify features that were both strongly associated with a given biome and present across multiple clades. Phyloglm is a generalized linear model that combines logistic and Poisson regression with a phylogenetic tree. Together, the intersection of RF model and phylogenetic regression analysis yielded 654 informative, phylogeny-independent features (adjusted p < 0.01, Gini score > 0.001; Fig. 2D, Fig. S1F).

**Figure 2.**
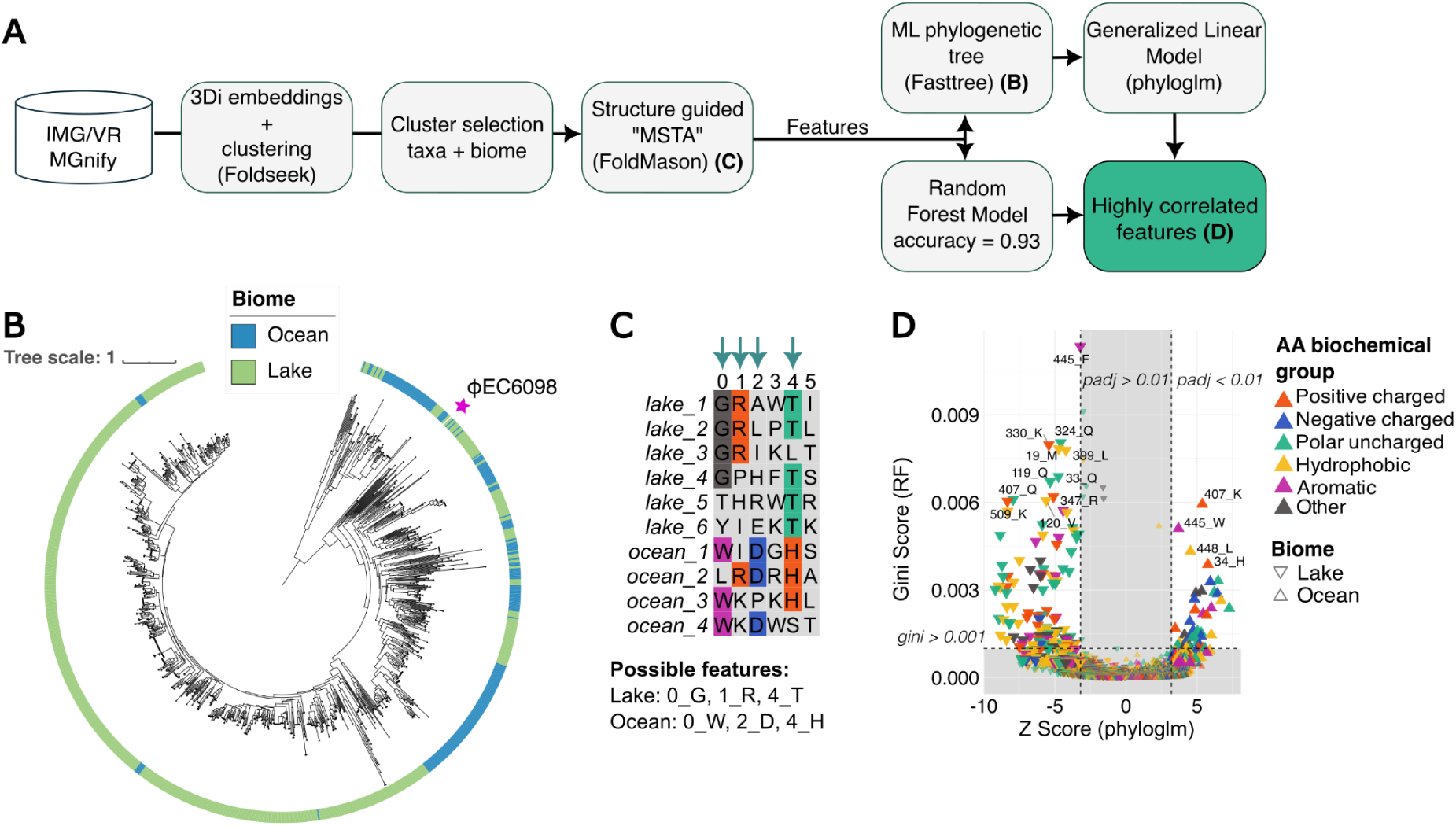
A) Workflow for identifying “Lake” or “Ocean” associated amino acid residues. For details see text. B) Trimmed phylogenetic tree of Gokushovirales MCPs (n=1,110) that were used in the random forest (RF) and phyloglm analyses. The purple star indicates the location of the ɸEC6098 reference protein. For the full tree, see Fig S3. C) A conceptual example of a multiple sequence alignment with arrows pointing to MSA positions that might be selected by either phyloglm (Ives and Garland, 2010) or the RF model (selected residues are colored). D) Distribution of biome-associated features. Positive and negative z-scores indicate features associated with the “Ocean” or “Lake” biomes, respectively. Features outside the grey area (phyloglm p_adj_ < 0.01, RF gini score > 0.001) were considered to be strongly associated. Features are coloured by biochemical amino acid properties.

Neither model picked up any informative features from a hypervariable region between positions 225-310 (Fig. S1E), which corresponds to the receptor binding domain, related to host specificity rather than environmental conditions^29^. Each feature was also labeled as either “lake associated” or “ocean associated” based on the phyloglm Z-score (Fig. 2D). Clusters of important features were distributed across the length of the protein (Fig. S1F). Specifically, clustered features were located on the protein “arm”, which reaches into adjacent pentamer, the intra-pentamer interface, the base of the receptor binding domain (RBD) and the external surface. Important features with specific amino acids such as Ser, Thr, Ile and Gly also clustered at specific regions on the protein (Fig. S5).

### Capsid surfaces and interfaces adapt differently to excess sodium ions

To investigate the structural and biochemical roles of the 654 “informative” features, we mapped them onto the capsid assembly of ɸEC6098 (pdb_00008des^30^) from the Protein Data Bank (PDB)^38^. ɸEC6098 is highly similar to the *Microviricetes* capsids used in this analysis (Fig. 2B), as shown by the placement of ɸEC6098 in a phylogenetic tree and the low root mean square deviation (RMSD) values between the ɸEC6098 MCP structure and the example structures visualized (Fig. S3, Table ST1).

Mapping feature importance scores (Gini scores) onto the ɸEC6098 structure revealed several residues that were associated with either high- or low-salinity environments (Fig. 3A). Residues with the highest Gini importance scores were concentrated at the base of the trimeric protrusion and the base of the arms reaching into the neighbouring pentamer (Fig. 3B). In the trimeric orientation these residues were observed at protein interfaces between adjacent pentamers (Fig. 3C). Further important features were also found on the external surface of the capsid (Fig. 3A).

**Figure 3.**
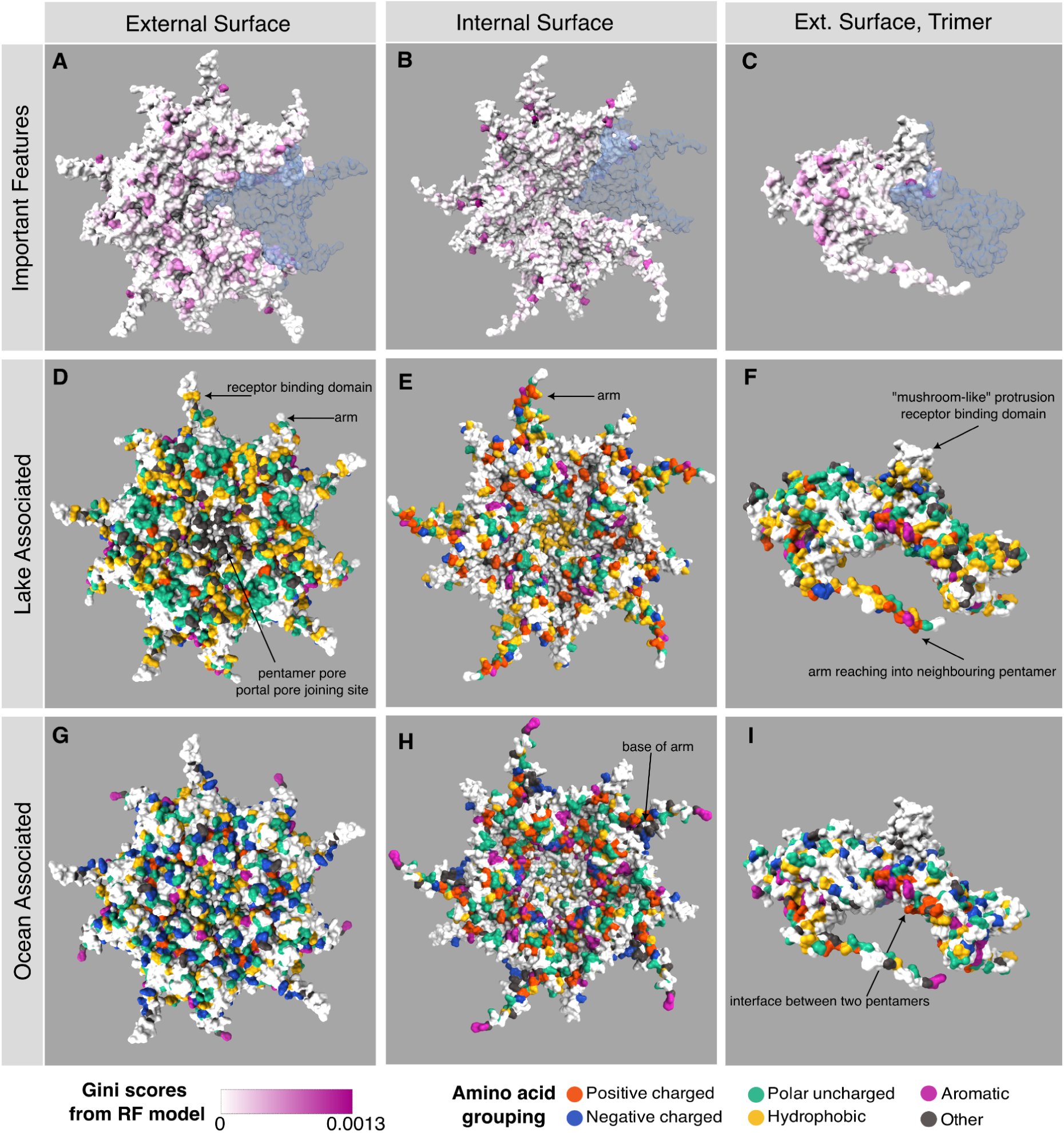
Informative features (n=654) mapped onto a reference capsid of enterogokushovirus ɸEC6098 (full protein length: 550 residues). A-C) Residues are coloured by the feature importance score from the RF model in the pentamer (A-B) and trimer (C) conformations. One of the monomers is coloured blue with 70% transparency to show the edges of pentamers and trimers. Further monomer-monomer boundaries are shown in Figure S4. D-F) Features strongly associated by the phyloglm model with the lake (“lake associated”, n=372) and (G-I) features strongly associated by the phyloglm model with the ocean (“ocean associated”, n=282) are coloured by groupings of biochemical properties of the amino acids.

We coloured the important residues by the amino acids corresponding to features that were either “lake associated” (Fig. 3D-F) or “ocean associated” (Fig. 3G-I). External surfaces of lake capsids were characterised by a combination of hydrophobic and polar uncharged residues (Fig. 3E), with an enrichment in Gly, particularly at the pore at the centre of the pentamer and across the external surface (arrow in Fig. 3D). Many lake-associated residues on the internal surface of the capsid were positively charged (Fig. 3E), likely reflecting capsid-stabilising interactions with the negatively charged ssDNA^39^. Indeed, previous nanoindentation studies have shown that full viral capsids are more resistant to indentation and breakage than empty capsids^40^. In freshwater, i.e. low ionic strength environments, capsids do not need electrostatic shielding of charge from the environment. In such environments, polar uncharged residues on the capsid exterior help maintain hydrophilicity and therefore solubility without adding net charge, and prevent the aggregation of viral particles. This may also explain why Gly, the smallest of all amino acids, highly flexible, and capable of forming tight loop structures, is favoured at the pentamer pore (center of the pentamer). The pore undergoes the largest conformational change by opening and serving as the extrusion site for the pilot protein through which DNA is injected at the beginning of an infection ^41^. These patterns can also be observed in Fig. S6 and S2 (purple triangles), which show closely grouped important features with Gly, Ser, and Thr enriched on the external surface of the majority of lake capsids.

External surfaces of ocean capsids were instead enriched in negatively charged residues (Glu and Asp) (Fig. 3G-I, S6). In saltwater, viruses face two challenges imposed by the high ionic strength of seawater. First, free ions can interfere with the charge-dependent electrostatic interactions that hold the capsid together, and second, saltwater can strip hydration from the capsid surface and promote aggregation. Therefore, an adaptation of amino acids on the capsid surfaces to seawater ions is expected to overcome these challenges. Indeed, previous studies have shown a similar enrichment of negative charge on the surface of halophilic proteins^9–12,42,43^. An interesting question that remained was why positively charged residues (Lys, Arg), which could equally pair with the abundant Cl⁻ in seawater, are not enriched instead. We suggest that the answer relates to charge density and hydration. In a high-ionic-strength solvent, water is drawn away from the protein surface to hydrate the abundant free ions (primarily Na⁺ and Cl⁻), causing dehydration of the capsid surface. Na⁺ has a high charge density and holds water tightly; carboxylate groups (Asp, Glu) are similarly high in charge density and strongly hydrated. This matched water affinity allows Na⁺ and carboxylate to form strong inner-sphere ion pairs without stripping away the carboxylate hydration shell, thereby keeping the surface hydrated and soluble. By contrast, the positively charged Lys and Arg and their seawater counterion Cl⁻ are relatively large and have low charge densities, allowing them to form weakly hydrated inner-sphere pairs. This weak bonding and the loose hydration shell around Cl⁻ would enable it to also form weak interactions with Lys and Arg residues in other protein subunits or other capsids, ultimately causing aggregation of the capsids rather than dispersal in the water column ^44^.

At the pentamer pore, glycine residues present in lake capsids were replaced by negatively charged, hydrophilic residues in ocean capsids, potentially indicating a more rigid interface for pilot protein binding in the ocean than in lake capsids. The presence of negative charge on the internal surface was unexpected, as many viral capsid interiors are typically positively charged so that interactions with the tightly packed, negatively charged ssDNA acts as an extra stabilisation mechanism^39,45^. Because microvirus capsids are likely semi-permeable^29^, seawater ions may enter the capsid and compete with the positively charged residues on the capsid interior for interactions with the ssDNA. We hypothesise that ocean capsids have evolved additional negative charge on their internal surfaces to sequester the cations, leaving DNA to bind instead with positively charged residues and maintaining capsid stability.

To complement these results, we also investigated which amino acids were overall surface-exposed or buried (Fig. S7). For this, we calculated the relative solvent accessibility (RSA) by normalizing the solvent-accessible surface area (SASA) from ChimeraX of each amino acid in each of the 1,110 proteins by their maximum theoretical solvent accessibility^46^. The maximum theoretical solvent accessibility was determined by Tien et. al. based on structures of ∼ 3000 proteins. This analysis confirmed that negatively charged residues Glu and Asp, polar residue Gln, and aromatic residue Trp were surface exposed in ocean capsids, but more buried in the lake capsids (Fig. S7, denoted by dashed line). Meanwhile, Arg, Ala, Ile, Phe, Cys and Pro residues are overall more buried in the ocean than in lake capsids (Fig. S7).

### Inter- and intra-pentamer interactions are enriched in hydrogen, cation-pi, and pi-pi bonds

When mapping the important features onto the ɸEC6098 capsid, we observed that several clustered at the base of the receptor binding domain (RBD) protrusion and at the interface between MCP subunits of adjacent pentamers. This suggested that inter-pentamer interactions, in addition to surface charge and capsid-DNA interactions, may also shift with the environment. To quantify these interactions, we extrapolated the interatomic contacts of a single monomer from the ɸEC6098 capsid structure in ChimeraX to all other proteins in the dataset via equivalent positions in the multiple sequence alignment, an approach justified by the high sequence and structural homology among all *Microviricetes* MCPs used in this study and the MCP of ɸEC6098 (Fig. S3, Table ST1). Because bond angles and true interatomic distances could not be translated by extrapolation, we adopted a conservative counting scheme which included only side-chain to side-chain and side-chain to backbone contacts, and with each amino acid pair represented by a single bond (Table 2).

Within monomers, hydrophobic interactions and hydrogen bonds together accounted for 60-80% of all contacts. Hydrophobic interactions dominated in both biomes, consistent with their central role in protein folding and packing (median 40% in lake capsids vs. 38% in ocean capsids), and H-bonds, which form between polar residues, were slightly more prevalent in ocean capsids (median 29% vs. 28%; Fig. 4A). We hypothesize that the substitution of hydrophobic surface residues for polar and acidic ones in ocean capsids is the reason for this difference.

**Figure 4.**
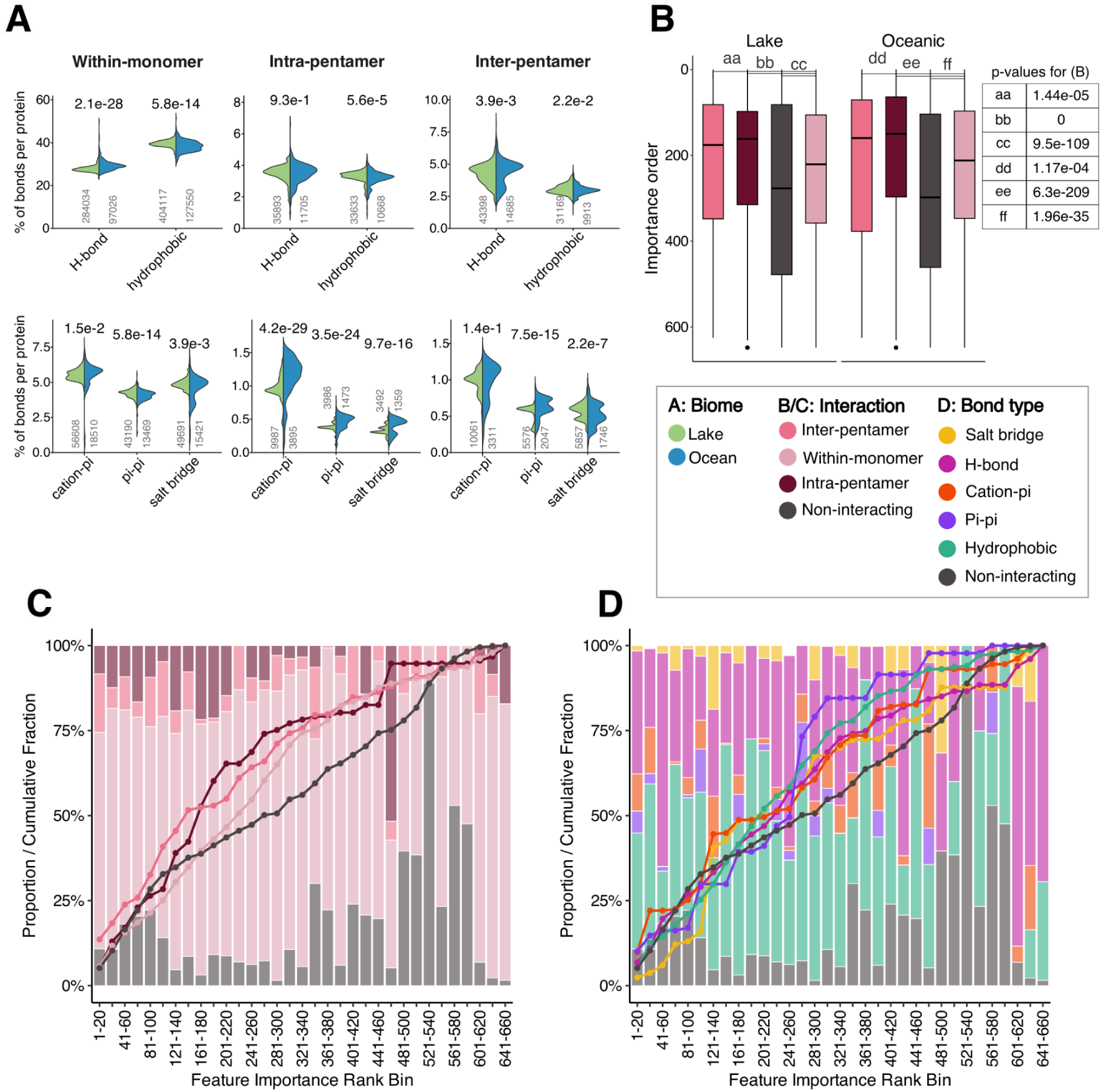
A) Percentage of residues with each bond type per protein at different monomer-monomer interfaces. Bond types are separated into two rows with different Y axis scales. Inter-pentamer bonds connect monomers outside the pentameric unit; intra-pentamer: bonds within the pentameric unit; and within-monomer: bonds formed within the monomer. Within-monomer bonds are excluded from the intra-pentamer count. The total number of bonds are shown under the x-axis in thousands. P-adjusted values are based on a Wilcoxon test (multiple testing correction by Benjamini-Hochberg). B) Importance rank of interacting residues. An importance rank of 0 corresponds to the most informative feature in the RF model. P-values are based on a Wilcoxon test. C) The percentage of interactions by interaction type within each importance rank bin (bars), where 100% corresponds to all features in that rank bin, and the cumulative percentage (line), where 100% corresponds to all the interactions of that type (e.g. 100% of inter-pentamer interactions). D) The percentage of interactions by biochemical bond type within each importance rank bin (bars) and the cumulative percentage (line). Table 2 shows which amino acids are involved in which biochemical bond types.

Within-monomer contacts in ocean capsids also showed slightly more cation-π interactions, which form between positively charged and aromatic residues, and slightly fewer π-π interactions and salt bridges, which form between aromatic residue pairs and oppositely charged residue pairs, respectively.

Considering interactions between MCP subunits, hydrogen bonds were more prevalent than hydrophobic interactions at both intra- and inter-pentamer interfaces, indicating a greater role for polar interactions in capsid assembly in both biomes (Fig. 4A). At intra-pentamer interfaces, cation-π, π-π, and salt bridge interactions were all significantly enriched in ocean capsids (Wilcoxon test, p = 2.8e-30, 6.92e-25, and 2.59e-16, respectively), while at inter-pentamer interfaces only π-π interactions were significantly enriched (p = 2.49e-15) and salt bridges were depleted (p = 1.2e-7). Salt bridges lose much of their stabilizing energy upon solvation and may be further disrupted by ions in the salt water, whereas cation-π interactions remain relatively resilient at solvent-exposed surfaces^42,43^. The depletion of salt bridges therefore suggests that some inter-pentameric interfaces may be more exposed or susceptible to destabilization in ocean capsids.

Ranking features by lake/ocean prediction importance showed that residues at intra- and inter-pentamer contacts were the most informative on average, followed by within-monomer contacts, with non-interacting residues the least informative (Fig. 4B). Half of all inter-pentamer bonds fell within the 150 most important features (Fig. 4C), and the most important of these clustered near the base of the RBD and the arm (Fig. S8), suggesting that adaptation at these interfaces is particularly important for maintaining structural stability across environments. Vihinen flexibility also drops sharply in this region in both biomes (Fig. S1, positions 175-225 and 310-350), indicating that rigidity at the base of the RBD is crucial for inter-pentameric stability and RBD function.

A second segment, between positions 440-450, also folds near the base of the RBD and contributes to inter-pentameric binding. Within it, F446W and F449L (lake to ocean) all ranked among the top 10 features in the random forest model, and constituted conservative substitutions with similar rigidity and hydrophobicity. One hypothesis for F446W is that the indole ring of Trp, which carries an N-H group capable of hydrogen bonding, reflects increased hydrogen bonding capacity at this position.

Finally, when ranked interactions were further resolved into specific bond types (Fig. 4D), the most notable feature was the large proportion of cation-π interactions among the top 40 features, reiterating the importance of aromatic residues and cation-π bonding in capsid stability.

### Ocean capsids are defined by specific amino acid substitutions outside of their biochemical groups

To test whether particular amino acids were consistently substituted for one another between lake and oceanic proteins, we examined the likelihood of finding a given amino acid in one biome at positions identified as important in the other (Fig. 5). The sequence logo in Figure 5A illustrates this approach. It shows how often glutamate (E, dark blue) appears as an important feature in the ocean biome, and how often glutamine (Q, teal) occupies those same positions in lake sequences.

**Figure 5.**
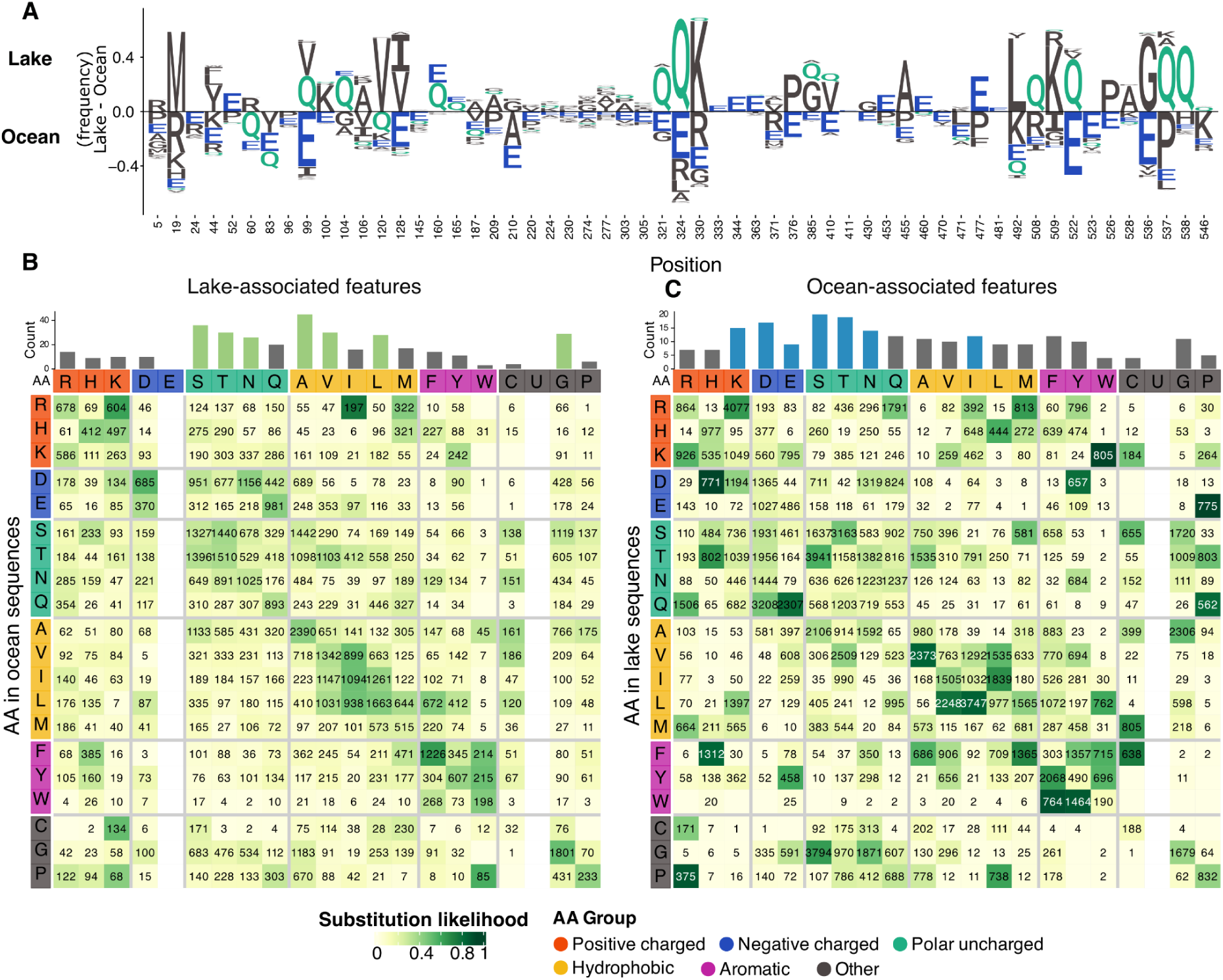
A) An example of preferred (reoccuring) amino acid substitutions shown through a sequence logo. Sequence logo shows all positions where a Glutamate (“E”) occurs in 5% or more of ocean sequences. B/C) Amino acid transitions at 653 important positions that distinguish Lake and Ocean sequences. Heatmap coloring shows the average likelihood of finding an amino acid in an ocean protein, given that the corresponding residue is predictive for the lake biome (B), or vice versa (C). For example, in (A), the likelihood of observing a “Q” in the lake sequences at positions with an E in the ocean sequences is higher than other amino acids. The numbers show how often this amino acid pair occurs between the ocean and lake protein sequences. The bar graph above the heatmap indicates the total number of times this amino acid occurs among the informative features for each biome. Colored bars are the amino acids with high counts. Note: There are no important lake-associated features containing Glu (E).

Charged polar residues were enriched among the top ocean-associated features, whereas uncharged polar and nonpolar residues dominated the lake-associated features (Fig. 5B/C), a pattern also visible when the features are mapped onto the protein structures (Fig. 3). Glutamate was entirely absent from the lake-associated features. The substitution heatmaps revealed an asymmetry in specificity between the two biomes (Fig. 5B-C). At the positions of significant lake-associated residues, the respective amino acids in ocean proteins had similar biochemical properties as their lake counterparts, reflected by darker coloured groups along the diagonal in the lake heatmap (Fig. 5B) (within-group). This suggests that substitutions in lake capsids are biochemically less specific than in ocean capsids (Fig. S9). In contrast, at ocean-associated positions, lake proteins often showed divergent substitutions to amino acids outside their biochemical group (between-group), as reflected by specific dark coloured squares outside of the diagonal (Fig. 5C). This suggests that adaptations to saline environments require not only presence of specific amino acids, for example Glu or Asp, but also absence of others, and that these substitutions are highly specific outside their biochemical group (seen in the long tail of the violins in Fig. S9 between-group panel). Specifically, 36 between-group substitution pairs had a likelihood of > 0.2 in ocean-associated features, compared with only 9 in lake-associated features, whereas within-group pairs above this threshold were similarly frequent in both (19 and 17, respectively) (Fig. S9).

From the ocean-associated features heatmap (Fig. 5C), we observed several between-group substitutions with high likelihoods, including Gln-to-Asp (0.25), Gln-to-Glu (0.4), Gly-to-Ser (0.33), Lys-to-Trp (0.49), and Glu-to-Pro (0.49), as well as recurring within-group substitutions such as Lys-to-Arg (0.29), Val-to-Ala (0.43), Leu-to-Val (0.4), Leu-to-Ile (0.44), Trp-to-Phe (0.46), and Trp-to-Tyr (0.45). The Gly-to-Ser and Gln-to-Asp/Glu substitutions frequently occurred on the outer surface of the capsid, contributing to a more polar and acidic outer surface (Fig. 3). The common within-group substitutions could be considered conservative, pointing to subtle shifts in biochemistry rather than large changes in charge or polarity. One example is the Leu-to-Ile substitution, which preserves hydrophobicity but increases rigidity at these positions, as Ile contains a beta-branched side chain that restricts backbone flexibility more than that of Leu. These substitutions were positioned between residues 20-100, where they occurred on either the internal surface or in buried positions (Fig. S6, black star).

Another example of conservative amino acid substitution was that of Arg-to-Lys. Despite both being positively charged, side chains of Lys and Arg behave differently in bonding. Lysine has a positive charge only at the end of its side chain, while in Arg it is delocalized over three nitrogens. This makes interactions with Lys more specific and directional, while Arg may interact with multiple residues simultaneously, promoting aggregation of capsids if located at the surface. A recent study showed that high Lys/Arg (K/R) ratios are strongly correlated with protein solubility in aqueous media^47^ and K/R ratios have been suggested for use in protein engineering as a measure of protein solubility^47,48^. Contrastingly, several studies have also reported a depletion of Lys from halophilic proteins^10,49,50^ due to a mutation of Lys-to-Glu common in halophilic proteins. In our analysis, Asp and Lys were both enriched on the surface of microvirus MCPs (Asp on the outer, and Lys on the inner surface) and substitutions between Lys and Glu were rare. Additionally, K/R ratios were significantly higher in ocean capsids than in lake capsids, both on the internal and external surfaces (Wilcoxon test, *p* = 9.55e-31 and *p* = 1.72e-42, respectively; Fig. S10), and Arg residues had significantly lower relative solvent accessibility in the ocean than in lake capsids (Wilcoxon test, *p* = 0.00016, Fig. S7). The discrepancy between the low K/R ratios reported for halophilic proteins and the high ratios we observed in ocean capsids might be explained by the fact that capsids are ultimately assembled and packaged inside the host cell. As intracellular salt concentrations can be either low or high depending on the haloadaptation strategy of the host, capsids may need to remain soluble and stable both inside the cell before lysis and under environmental conditions after release.

## Conclusion

In this study, we combined metagenomic sequence mining, machine learning, and structural analyses to identify molecular signatures of salinity adaptation in microvirus capsids. Ocean capsids were more acidic, hydrophilic, and flexible than freshwater ones, a signature captured by a random forest model with 93% balanced accuracy. Mapping 653 biome-associated features onto the capsid structure revealed enriched negative charge on both external and internal surfaces, which we hypothesize reflects hydration under high salinity and cation sequestration within the capsid. Despite increased overall flexibility, ocean capsids exhibited localised rigidity at the pentamer pore and receptor-binding domain. Their monomer interfaces also displayed distinct bonds, including increased cation–π and π–π interactions and fewer salt bridges. Substitution patterns further suggested that adaptation to the ocean ecosystem imposes more specific residue requirements than freshwater adaptation, and that haloadaptation strategies may be different for proteins that need to remain stable both inside and outside the cell.

As our pipeline (XtrAActor) requires only protein/genomic sequences and biome metadata as input, it offers an extendable framework for investigating the adaptations of any kind of protein to environmental conditions, such as salinity, temperature, pH, or other phenotypes. We have made this pipeline publicly available in a Github repository for researchers to investigate the adaptations of their proteins of interest. By linking sequence variation to structural features, our approach generates specific hypotheses about the mechanisms underlying environmental adaptation and reveals how environmental selection shapes protein evolution, viral stability, and persistence across ecosystems. Furthermore, by learning from existing metagenomic datasets of natural systems, our approach could guide the engineering of more robust viral capsids or proteins for biotechnological applications, including those requiring resistance to extreme conditions.

## Methods

Scripts for all data retrieval, filtering, and visualization, as well as various analyses are available in the following repository: https://github.com/vmkhot/RF-model-nextflow/tree/main/protein_analyses. Additionally, a nextflow pipeline was built to cluster sequences, create a multiple sequence alignment, generate random forest models, and output Gini importance scores for the features https://github.com/MGXlab/XtrAActor. This pipeline only requires biome metadata and genome/protein sequences as input and can be generalized to any protein.

### Major Capsid Protein Sequence Retrieval and Filtering

Metagenomic protein sequences were downloaded from IMG/VR v4.7 ^20^ and MGnify (v2023) ^21^. These were filtered for ‘major head proteins’ using Pharokka ^51^, which included 22 PHROG ^24^ categories with the gene product as “major head protein”, yielding a total of 1.4 million major capsid proteins (MCPs). We focused our analysis on phages from the family *Microviricetes* by only including PHROG 514 MCPs. These were considered to be MCPs with the typical single-jelly-roll fold, and were used for subsequent analyses in this manuscript. Microvirus MCPs were further filtered for a sequence length between 250-600aa, and for those which had biome metadata categories of either “Environmental;Aquatic;Freshwater;Lake” or “Environmental;Aquatic;Marine;Oceanic” biomes (14). This resulted in 1,338 “Lake” and “Oceanic” proteins.

To visualize the sequence similarity of all major capsid proteins, they were then clustered using diamond linclust^53^ with the parameters “--approx-id 70” and “--member-cover 80”. Cluster representative were then selected using seqtk r133^54^ and aligned to each other using diamond blastp^22^ with the parameters “-k 100”, “--id 50”, “--evalue 0.00001”, “--query-cover 80”, “--subject-cover 80”, “--outfmt 6 qseqid sseqid pident length mismatch gapopen”. Blastp results were further filtered to remove self-hits before creating a sequence similarity network using the -log(evalue) as edge weights.

### Structural Clustering

To group together sequences from homologous proteins for improved multiple sequence alignments, *Microviricetes* capsid proteins were clustered based on structural similarity using Foldseek v10-941cd33 ^31^, by first creating a database of 3Di embeddings using ProstT5 model prostt5_10_941_weights ^55^ and then clustering using Foldseek at 50% identity and 70% bidirectional coverage.

Only sequences from clusters with similar capsid morphology to the ɸEC6098 *Enterogokushovirus* ^30^ were retained, resulting in 1,110 proteins for downstream analysis: 824 from the “Lake” biome and 286 from the “Oceanic” biome. This was to ensure that multiple sequence and structural alignments to the ɸEC6098 capsid sequence and assembly would be as close as possible, and to exclude PhiX type capsids with spurious biome annotations from the dataset. DNA from ɸX174 microviruses is commonly used as a spike-in for quality control in Illumina sequencing, but are often not filtered out from the data. This may confound downstream analyses, including inferring associations between these viruses and biome labels in the case of metagenomic sequencing.

Protein properties, namely molecular weight, isoelectric point, charge at pH 7, aromaticity, flexibility, and grand average of hydropathy (GRAVY), of “Lake” and “Ocean” microvirus capsid sequences were then calculated with biopython v1.6 ^23^.

### Multiple Sequence and Structure Alignment

Retained proteins with their 3Di embeddings were aligned using FoldMason ^56^ with default parameters, which employs a structure-guided multiple sequence alignment (MSTA) algorithm. To improve alignment quality, all sequences were additionally aligned to the major capsid protein of ɸEC6098 as a reference using mafft ^57^ with the “--add-sequences” parameter.

### Random Forest Classification

Features representing a position and amino acid (e.g. 0_G, 0_A etc.) were extracted from the MSTA by one-hot-encoding the alignment. To find specific features that associated with either a “Lake” or “Ocean” biome, alignment-derived features were then used to train a random forest model from the scikit-learn ^58^ package in Python “RandomForestClassifier()”, with a train-test split of 80/20. The optimal fit parameters were selected by hyperparameter tuning using RandomizedSearchCV, which resulted in the following parameters being chosen: “class_weight=’balanced’, max_features=’log2’, min_samples_split=5, n_estimators=1000”. Model performance was evaluated using balanced accuracy.

### Phylogeny-aware Generalised Linear Modelling

As features may be correlated to a biome due to shared ancestry, we also used a generalized linear model optimized to overcome phylogenetic bias, named phyloglm ^37,59^, in R, in addition to the random forest model.

For this analysis, we first produced a phylogenetic tree of the 1,100 *Gokushovirales (Microviricetes)* capsid sequences using the multiple sequence alignment from the previous step and FastTree v2.0^60^ using default parameters. The inputs to the phyloglm model included the features, biome-associated metadata for the capsid sequences and the phylogenetic tree. The R script used for phyloglm can be found here: https://github.com/vmkhot/labjournal/tree/main/phyloglm.

All features were finally ranked by both Gini importance scores from the random forest model as well as z-scores from phyloglm. Features meeting thresholds of adjusted p < 0.01 and Gini score > 0.001 were considered both informative and phylogenetically un-biased, yielding 654 features for subsequent analyses.

### Mapping features onto ɸEC6098 capsid assembly

The ɸEC6098 capsid assembly (pdb_00008des) ^30^ was used extensively in this manuscript as a reference protein/capsid. Using ChimeraX ^61^, we mapped Gini importance scores from 654 features onto this capsid assembly to visualize where on the capsid features selected as important were localized. These features were coloured by Gini score and biochemical properties as shown in Table 1. Scripts for mapping features onto proteins structures in ChimeraX are here : https://github.com/vmkhot/RF-model-nextflow/tree/main/protein_analyses/chimera_scripts

**Table 1:** Biochemical groupings of amino acids as defined in this study.

| Biochemical grouping | Amino acids |
|---|---|
| Positively charged | Arginine (Arg, R), Histidine (His, H), Lysine (Lys, K) |
| Negatively charged | Aspartate(Asp, D), Glutamate (Glu, E) |
| Polar uncharged | Serine (Ser, S), Threonine (Thr, T), Asparagine (Asn, N), Glutamine (Gln, Q) |
| Hydrophobic | Alanine (Ala, A), Valine (Val, V), Isoleucine (Ile, I), Leucine (Leu, L), Methionine (Met, M) |
| Aromatic | Phenylalanine (Phe, F), Tyrosine (Tyr, Y), Tryptophan (Trp, W) |
| Other | Cysteine (Cys, C), Glycine (Gly, G), Proline (Pro, P) |

#### Contacts

From the ɸEC6098 capsid assembly structure in ChimeraX, we also calculated the residue-residue contacts for each position, with a maximum distance of 6Å, using the “contacts” command. We then extrapolated to other MCPs via the original multiple sequence alignment as follows. Each position on the ɸEC6098 MCP was mapped to an equivalent position in protein X (example protein) via the original MSTA and the amino acid was changed to corresponding amino acid at that position in protein X. Contacts and bond types were then inferred for protein X within the capsid assembly if the distance to another residue was ⩽ 5.5 Å and the amino acids could form a bond based on their biochemical properties (Table 2). If protein X contains additional residues that cannot be mapped to ɸEC6098 MCP, then it is not used.

**Table 2.** Maximum distances allowed between amino acid side-chains, and the possible amino acids which are involved in each bond type. Arrow denotes amino acids which will form bonds: ↔

| **Bond type** | **Maximum distance (Å)** | **Acceptable amino acids** |
|---|---|---|
| Hydrogen | 3.5 | Ser, Thr, Tyr, Asn, Gln, His, Lys, Arg, Asp, Glu, Trp, Cys |
| Salt bridge | 4.0 | Asp, Glu (negative) ↔ Lys, Arg, His (positive) |
| Cation-pi | 5.5 | Lys, Arg (cation) ↔ Phe, Tyr, Trp |
| pi-pi (aromatic ring stacking) | 5.5 | Phe, Tyr, Trp |
| Disulfide bridges | 4.5 | Cys ↔ Cys |
| Hydrophobic interaction | 4.5 | Ala, Val, Leu, Ile, Met, Phe, Trp, Tyr, Pro |
| Van der Waals interaction | 4.5 | All amino acids |

As atomic bonds and interactions are dependent on the side-chain angles and geometry, in addition to the distance, we took a highly conservative approach by collapsing atom-level bonds to single residue-residue contacts with either side-chain-side-chain or side-chain-backbone contact within defined distance thresholds (Table 2). Each residue-residue pair was represented by its closest contact only based on a hierarchy of bonds: disulfide bridge, salt bridge, cation-pi, pi-pi, hydrogen bond and hydrophobic interaction. E.g. If a residue pair can make both a salt bridge and a hydrogen bond, the contact is classified as a salt bridge. This likely underestimates the total number of bonds, but we expect the overall trends to be preserved. Contacts were further classified into four categories based on their position in the capsid assembly: “self”: contacts within the same monomer, “intra-pentamer”: contacts between monomers at the 5-fold axis, “inter-pentamer”: contacts between monomers outside the 5-fold axis, and “non-interacting”: residues forming no contacts.

#### Solvent accessible surface area (SASA) calculations

Each of the microvirus capsid proteins used for model training (n=1,110) was folded using ColabFold^62^. Solvent-accessible surface area (SASA) was then calculated for the top-ranked folded model of each sequence using a custom ChimeraX script (available at: https://github.com/vmkhot/RF-model-nextflow/tree/main/protein_analyses/chimera_scripts). As capsid proteins assemble into a multimeric shell rather than existing as isolated monomers, SASA calculated from the monomeric structure alone overestimates true surface exposure. To correct for this, we applied a reduction factor of 0.54, derived from the difference in SASA between the assembled ɸEC6098 capsid and its isolated monomer, to approximate each protein’s SASA in its capsid-bound conformation. Relative surface area (RSA) was then calculated for each residue by normalizing the corrected SASA to its theoretical maximum value, as defined in Tien et al. 2023^46^.

### Amino acid substitutions analysis

To examine whether certain amino acid substitutions occur more frequently than others between the lake and ocean biomes, we calculated the average substitution likelihood for each lake- or ocean-associated feature (654 important features). For every important feature, we calculated the likelihood of observing each possible amino acid at the same position in the opposite biome. The script used for this calculation is available in the paper repository: https://github.com/vmkhot/RF-model-nextflow/tree/main/protein_analyses/.

## Acknowledgements

The authors would like to thank Malte Siemers, José Lopez, and Maria Beatriz Walter Costa for helpful discussions during data analysis, and Aristeidis Litos for his feedback on the manuscript.

## AI Statement

The AI agents Claude v5.5 by Anthropic and ChatGPT v4 and v4.5 were used in this study to make programming scripts for data analysis and editing the manuscript text.

