## Supplementary Figures and Tables for "Structural signatures of salinity adaptation in *Microviricetes* capsids"

### Supplementary Material

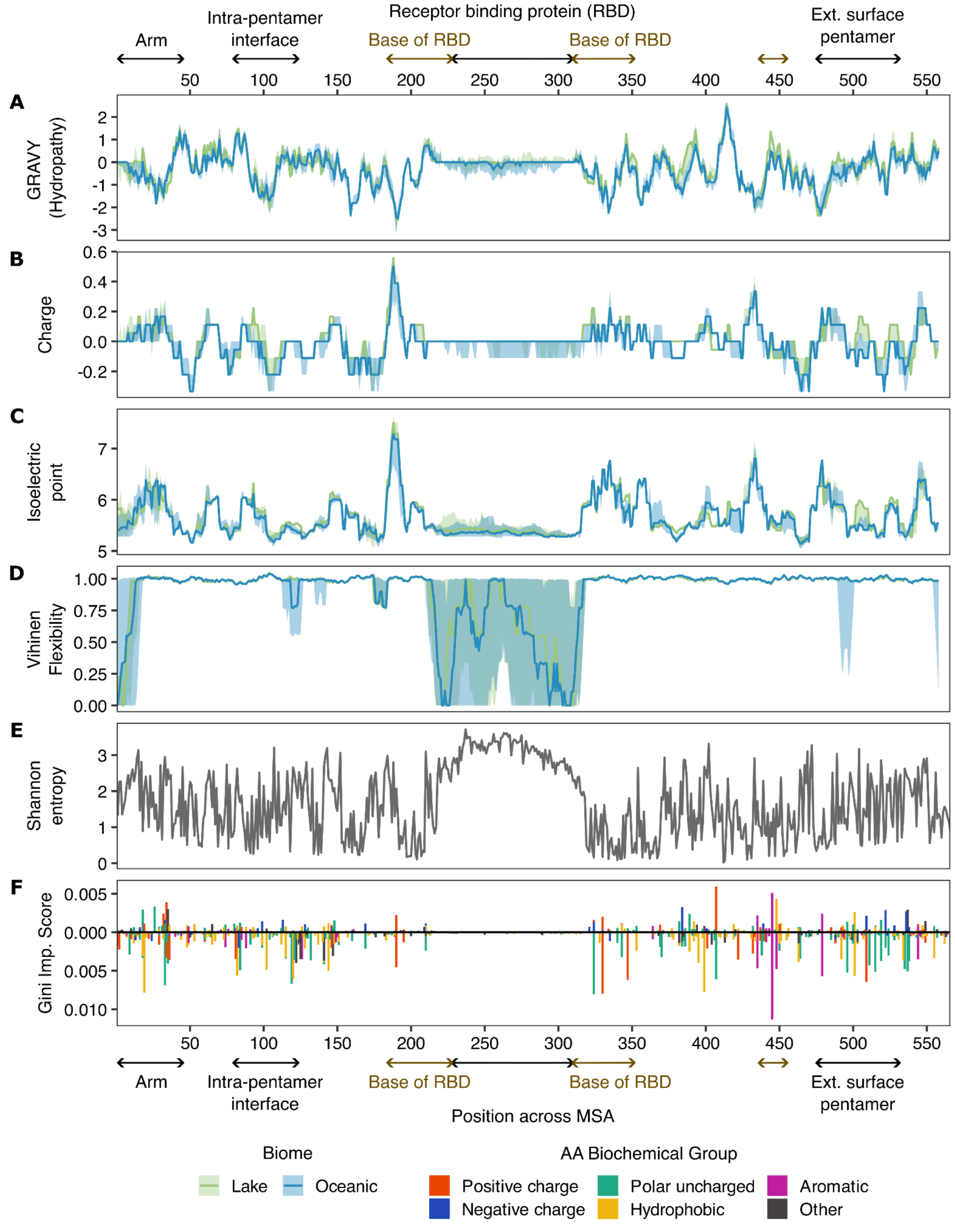


***Figure S1.*** *Protein properties, GRAVY index (A), charge (B), isoelectric point (C), and flexibility (D), shown along the length of the multiple sequence alignment (MSA), coloured by biome. The solid line shows the median value at each MSA position (sliding window: 9 amino acids). The shaded area shows 1 standard deviation. E) Shannon entropy calculated from all sequences in the MSA. F) Important features (n=654) as selected by the RF model and phyloglm shown along the MSA, coloured by the biochemical group of the amino acid in each feature.*

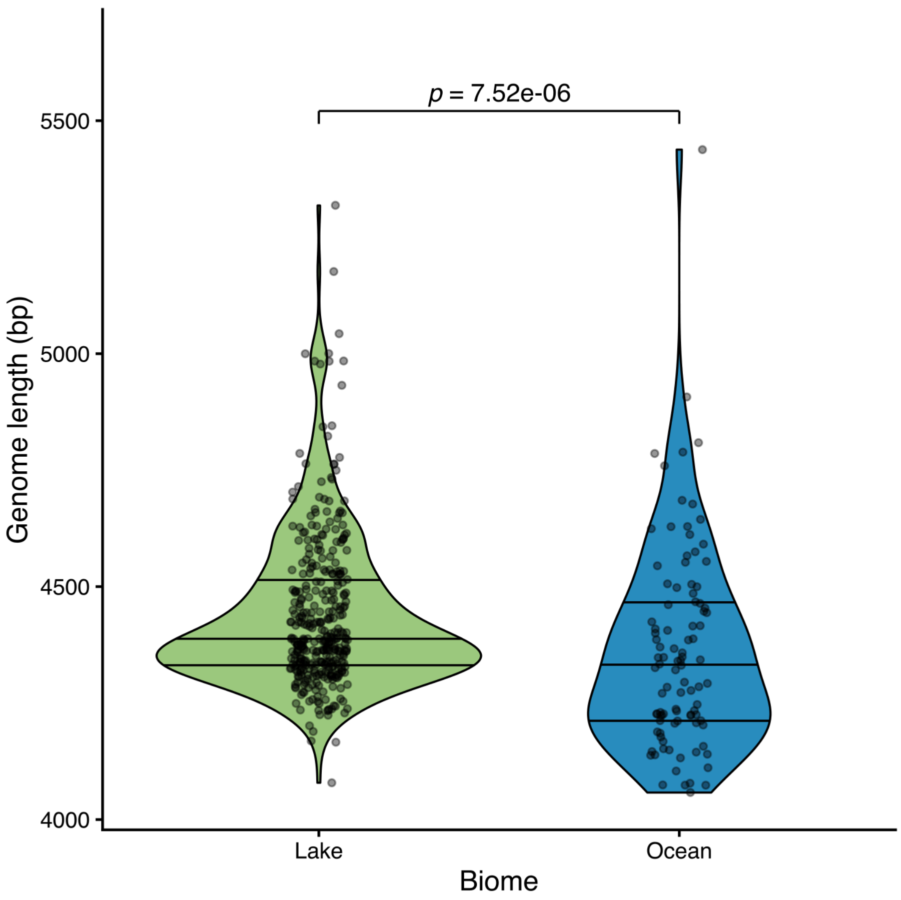


***Figure S2:*** *Genome length vs. biome of microviruses from IMG/VR used in this study. Full length microvirus genomes were selected for this comparison using CheckV* [1]*. Mann-Whitney test was used to compare genome length between the biomes.*

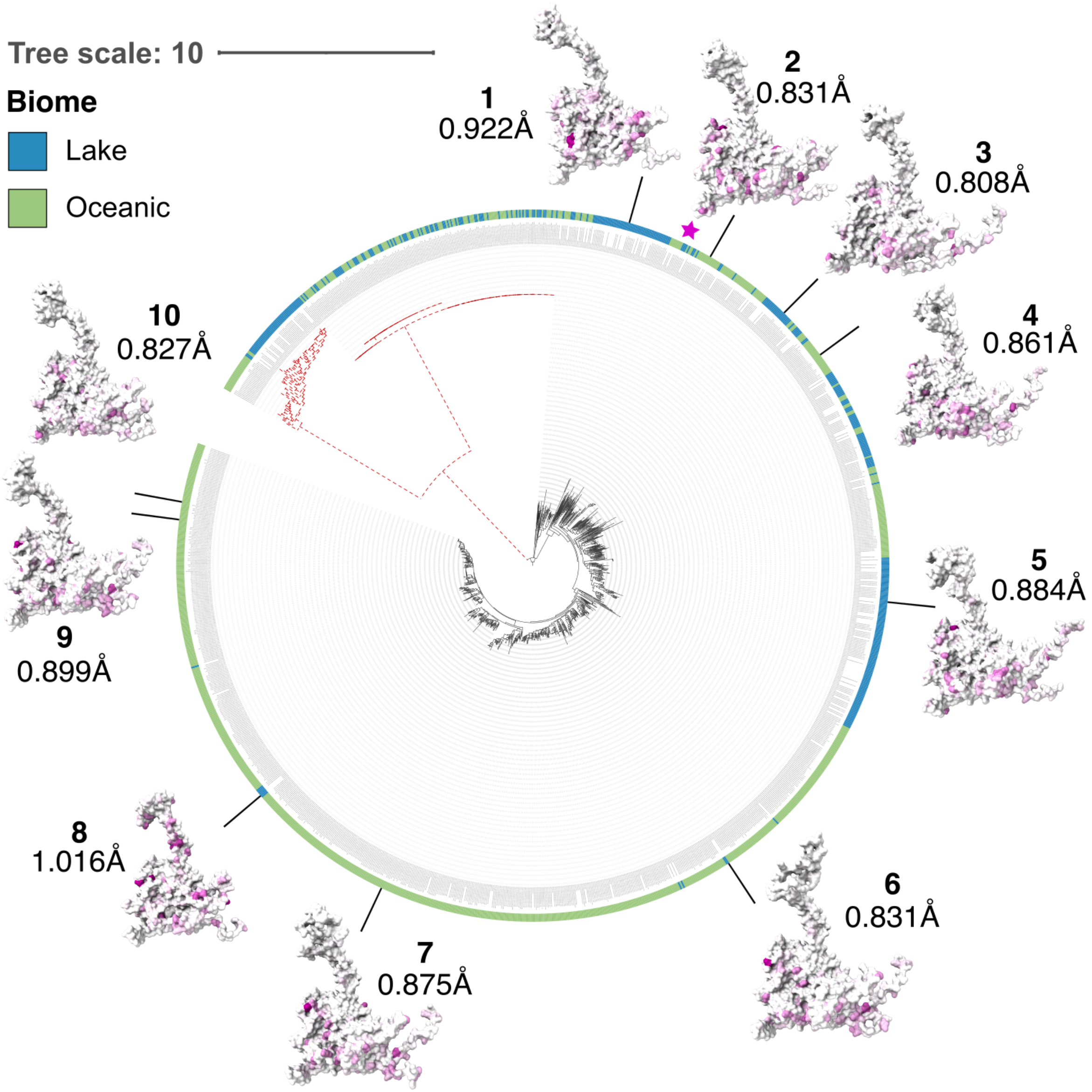


***Figure S3.*** *A phylogenetic tree based on* Microviricetes *capsid proteins. The tree was created using a multiple sequence alignment by Mafft followed by FasttreeMP[2,3], both with default parameters. Proteins on the red-coloured dashed branches were removed from the analysis as they produced a capsid morphology similar to ϕX174 rather than ϕEC6098. Placement of reference protein ϕEC6098 (8DES) is highlighted with a star. Examples of proteins from around the tree were visualized, and their corresponding root mean square deviation (RMSD) values to ϕEC6098 are shown. For details on these cases, see Table ST1.*

***Supplementary Table ST1:*** *Example proteins, their metadata, and their structural alignment to ϕEC6098 shown in angstroms. Proteins are visualized in Fig. S1. RMSD: root mean square deviation.*

| Number in Figure S1 | Protein name | Biome | Database | Fraction of pruned pairs out of total | RMSD to EC6098 (Å) |
|---|---|---|---|---|---|
| 1 | MGYP002641143952 | Ocean | MGnify | 293/400 | 0.922 |
| 2 | IMGVR_UViG_3300020042_009324_3300020042_Ga0206640_10135673 | Lake | IMG/VR v4.7 | 398/470 | 0.831 |
| 3 | MGYP003333885128 | Ocean | Mgnify | 366/423 | 0.808 |
| 4 | IMGVR_UViG_3300012282_000044_3300012282_Ga0157136_10002142 | Lake | IMG/VR v4.7 | 410/465 | 0.861 |
| 5 | MGYP001198978755 | Ocean | Mgnify | 406/462 | 0.884 |
| 6 | MGYP003704062755 | Ocean | Mgnify | 414/470 | 0.831 |
| 7 | IMGVR_UViG_3300043786_000143_3300043786_Ga0453106_0029613 | Lake | IMG/VR v4.7 | 408/467 | 0.875 |
| 8 | IMGVR_UViG_3300009526_000008_3300009526_Ga0115004_100234614 | Ocean | IMG/VR v4.7 | 269/372 | 1.016 |
| 9 | IMGVR_UViG_3300027697_000815_3300027697_Ga0209033_10097072 | Lake | IMG/VR v4.7 | 412/470 | 0.899 |
| 10 | IMGVR_UViG_3300012758_000284_3300012758_Ga0138285_11887812 | Lake | IMG/VR v4.7 | 410/454 | 0.827 |

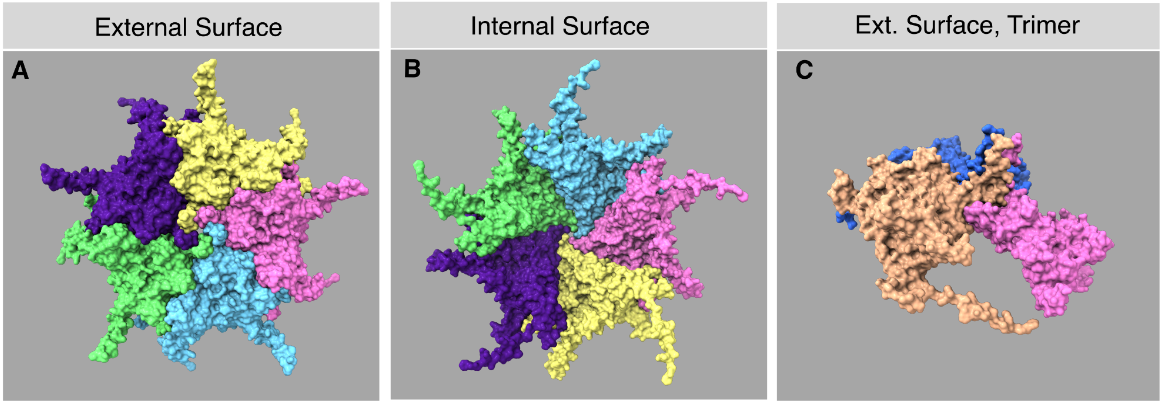


***Figure S4.*** *Outer (A) and inner surfaces (B) of the ϕEC6098 reference assembly pentamer and trimer (C) coloured by different subunits to show the boundaries between the monomers*.

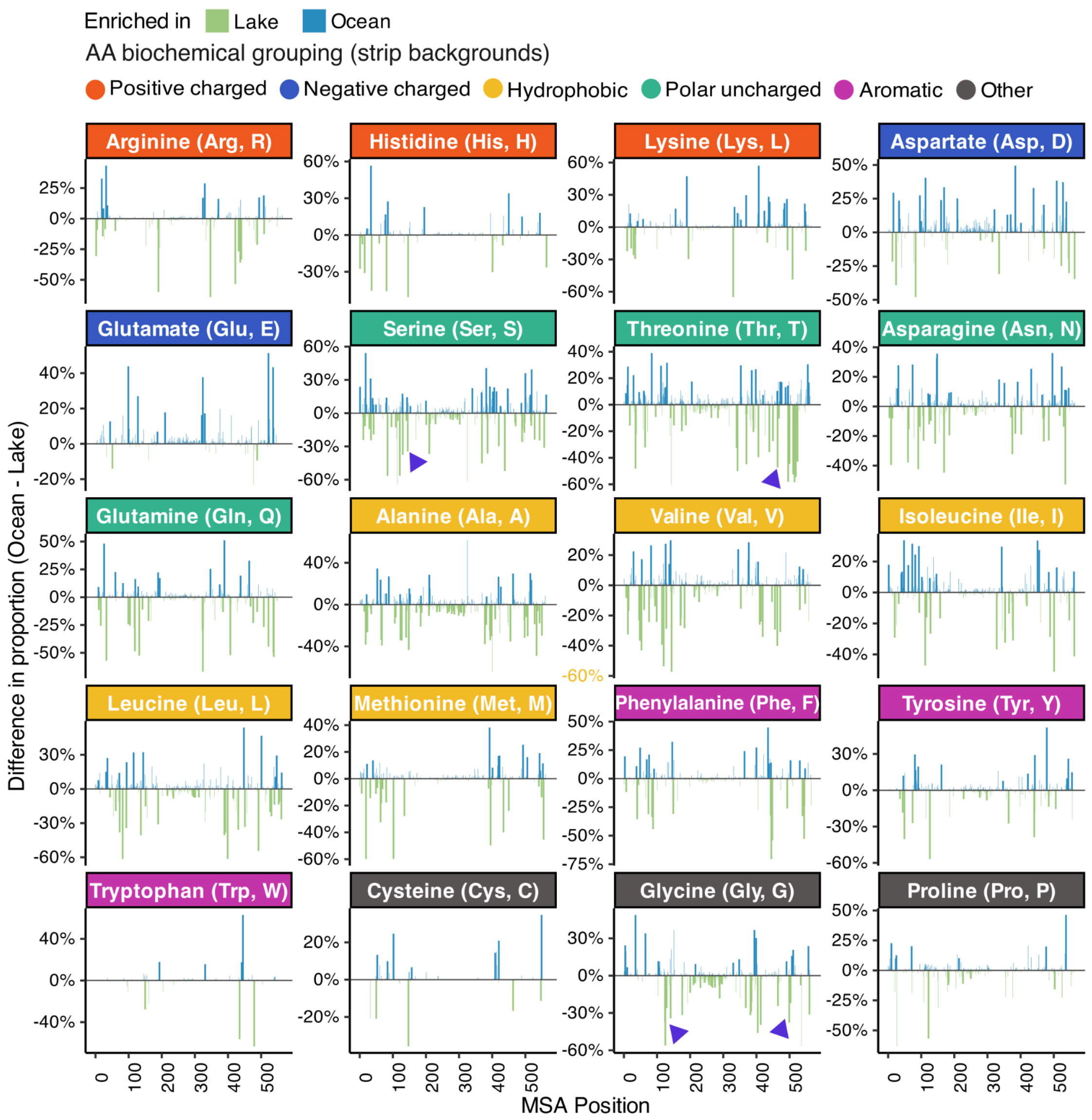


***Figure S5.*** *Difference in the number of Ocean and Lake sequences (Ocean - Lake) containing each amino acid across the MSA. Wider bars indicate positions and amino acids that are important features. Subplot headers are coloured by biochemical grouping of the amino acids.*

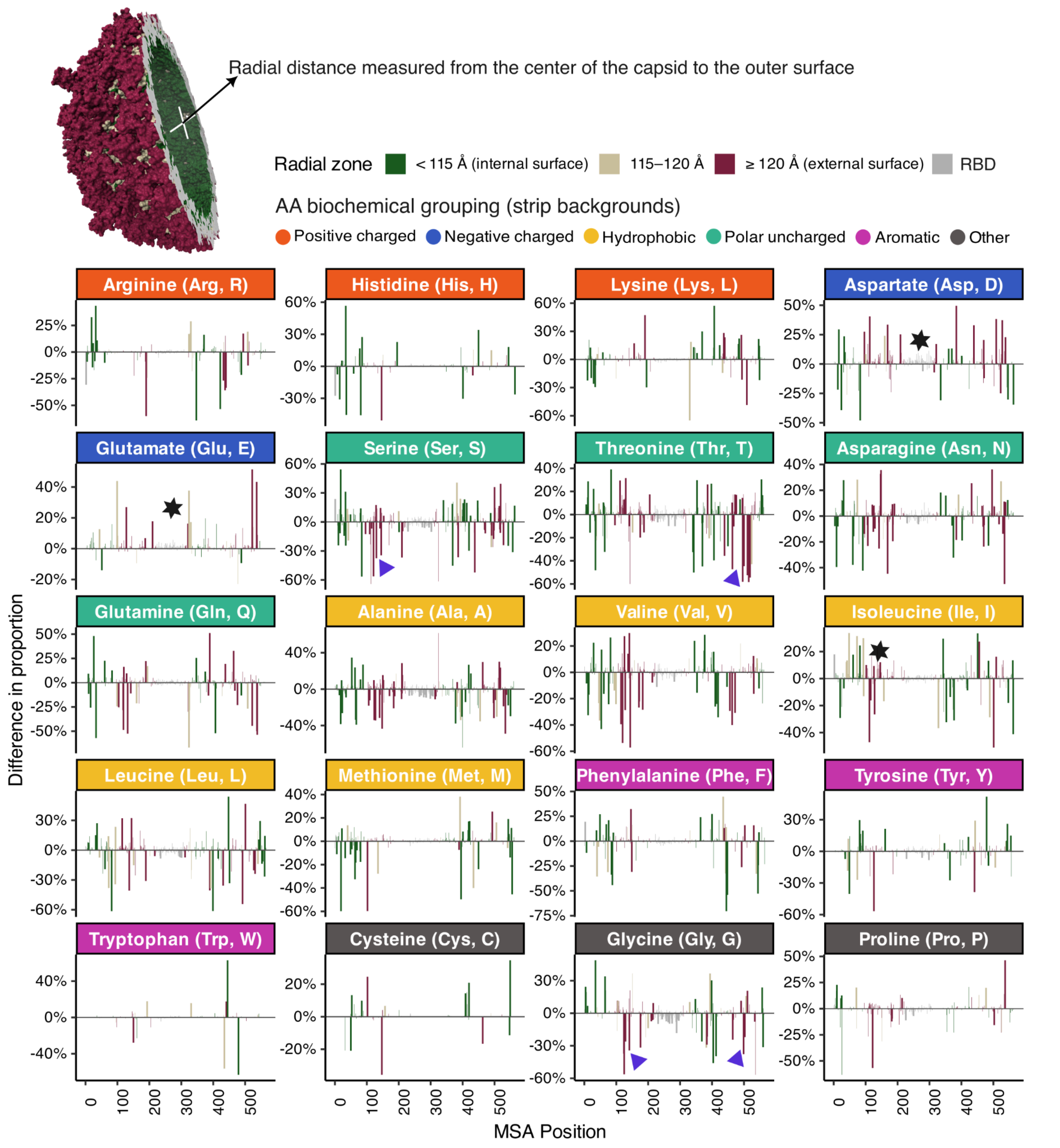


***Figure S6.*** *Difference in the number of Ocean and Lake sequences (Ocean - Lake) containing each amino acid across the MSA. Bar colours indicate whether the amino acid is exposed on the internal surface (green), external surface (maroon), or is buried (beige) (also shown by the capsid hemisphere). Wider bars indicate positions and amino acids that are important features. Black stars highlight some amino acids of interest discussed in the text.*

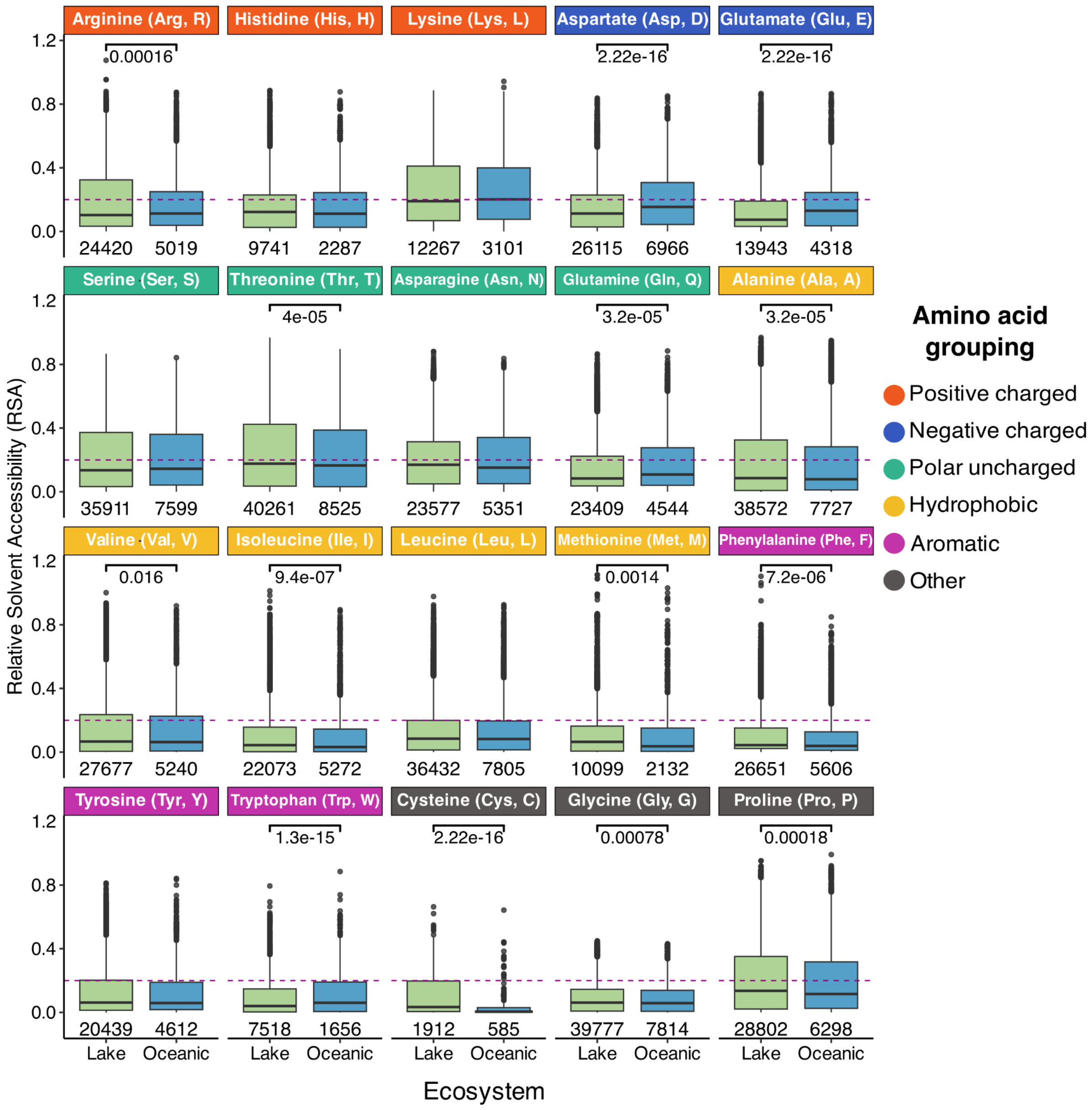


***Figure S7.*** *Relative solvent accessibility of each amino acid in Lake and Ocean proteins. RSA was calculated by calculating the solvent-accessible surface area (SASA) for each protein in its monomeric form, reducing the SASA using a factor calculated from the reference protein, and then dividing by the maximum solvent accessibility as described by Tien at al.*[4]*. Residues with RSA < 0.2 are buried residues denoted by the dashed line (compared to surface-exposed). P-values were calculated using the Wilcoxon test.*

Important features at the inter-pentameric interface

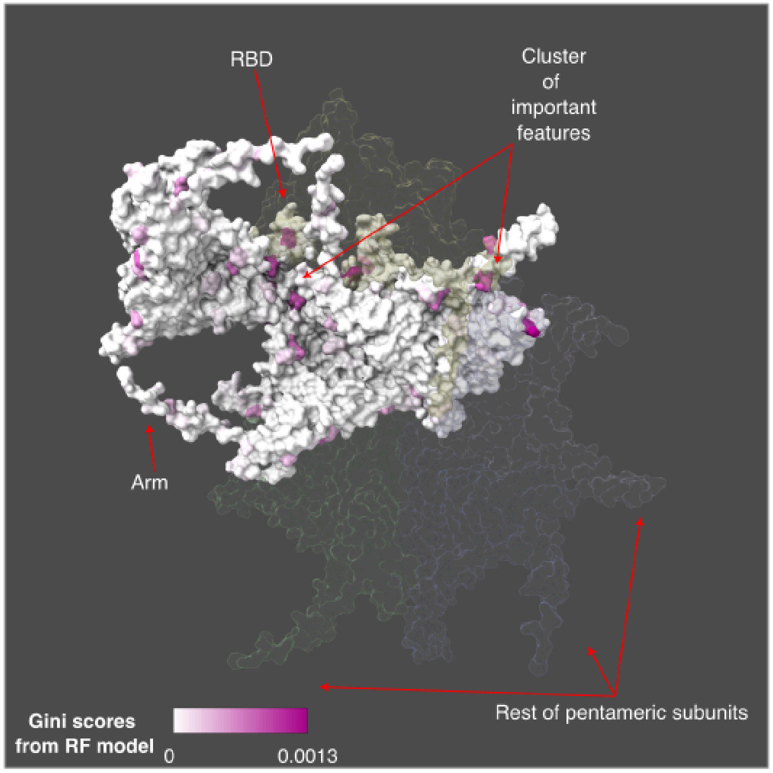


Important features at the intra-pentameric interface

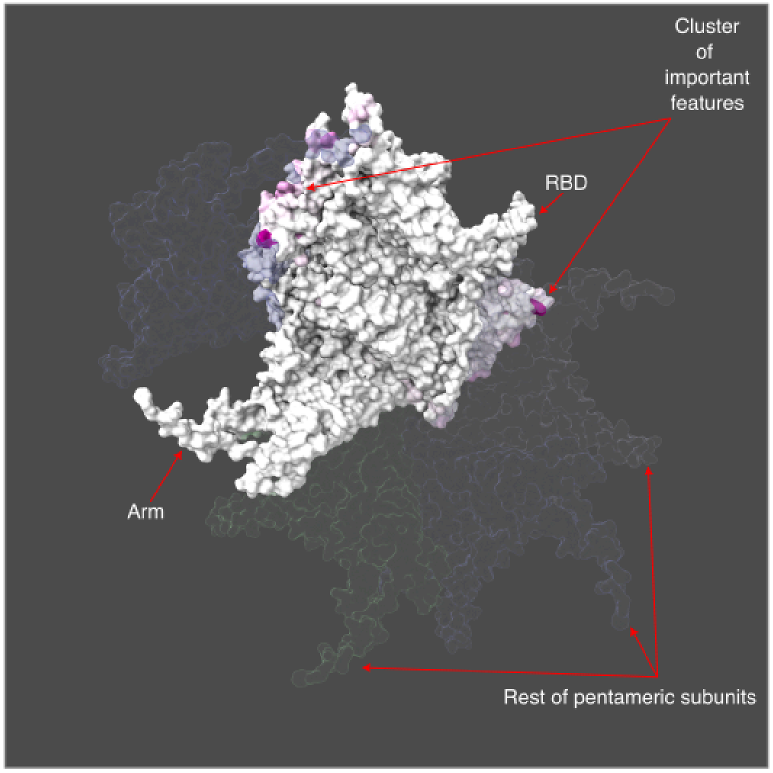

***Figure S8:*** *Important features at inter (top) and intra (bottom) -pentameric interfaces. Only important features at the interfaces are shown defined by residues which contact residues in another monomer. RBD: receptor binding domain*

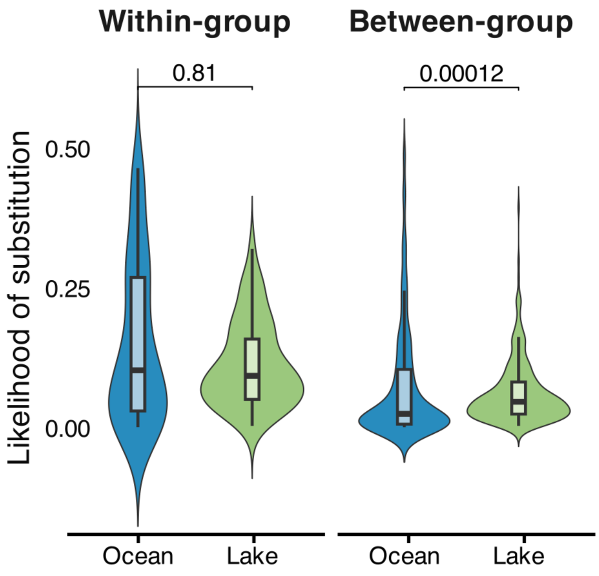


***Figure S9:*** *Likelihood of amino acids substitutions classified by within-group or between-group based on biochemical groups and which biome it is an important feature in.*

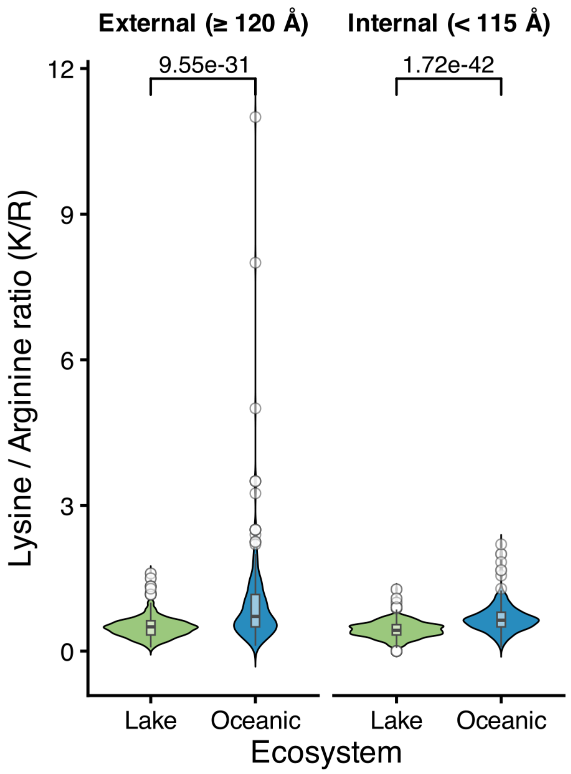


*Figure S10: Lysine to Arginine ratios on the internal and external surfaces of capsid proteins*
